# The stepwise endonuclease activity of a thermophilic Argonaute protein

**DOI:** 10.1101/821280

**Authors:** Guanhua Xun, Qian Liu, Yuesheng Chong, Zhonglei Li, Xiang Guo, Yinhua Li, He Fei, Kai Li, Yan Feng

**Affiliations:** State Key Laboratory of Microbial Metabolism, School of Life Sciences and Biotechnology, Shanghai Jiao Tong University, Shanghai 200240, P. R. China; Department of Obstetrics and Gynaecology, The Fifth People’s Hospital of Shanghai Affiliated to Fudan University, Shanghai 200240, P. R. China; GeneTalks Biotechnology Inc., Changsha, Hunan 410013, China

## Abstract

Thermophilic Argonaute proteins (Agos) can function as endonucleases via specific guide-target base-pairing cleavage for host defense. The ability to cleave target DNA sequences at any arbitrary sites endows them with reprogramed DNA capacity. Here, we identify that an Ago from the hyperthermophilic archaeon *Pyrococcus furiosus* (*Pf*Ago) shows a stepwise endonuclease activity, which is demonstrated by the double strand DNA cleavage directed by a single guide DNA rather than canonical one pair of guide DNAs. We reveal that the cleavage products with 5’-phosphorylated ends can used as the renewed guide which is capable to induce next round of cleavage to complementary sequences of target DNA. By combining the *Pf*Ago stepwise endonuclease activity followed by target DNA amplification, we establish a rapid and specific platform for the unambiguously multiplex gene detection, termed RADAR (Renewed-gDNA Assisted DNA-cleavage by Argonaute). In the end, RADAR was applied to distinguish of human papillomavirus of serotypes in patient samples in a single reaction, suggesting that our technique would be adopted for diagnosing application.

## Main Text

Argonaute proteins (Agos) play important roles in a wide range of biological processes including gene regulation and host defense via interacting with nucleic acids molecules [1-4]. Prokaryotic Agos are more diverse in their biochemical behavior than their eukaryotic counterparts. Thermophilic Agos have attracted increasing interest due to their unique endonuclease activity as directed by guide DNA/RNA molecules [5-8]. An Ago from the hyperthermophilic archaeon *Pyrococcus furiosus* (*Pf*Ago) can perform precise DNA cleavage directed by small 5’-phosphorylated single strand DNA (ssDNA) as guide DNA (gDNA) at 95 °C [7, 9]. The canonical cleavage product of *Pf*Ago was observed with cleavage occurring opposite nucleotide 10/11 of the gDNA. The thermophilic Agos from *Thermus thermophilus* (*Tt*Ago) and *Methanocaldococcus jannaschii* (*Mj*Ago) show the similar endonuclease activity to *Pf*Ago, but differs in the substrate spectra or optimal temperature [8, 10]. Moreover, it is noticed that double strand DNA (dsDNA) substrates could be chopped upon long time incubation at high temperature under a guide-free condition [11, 12], which indicates that the ssDNA generated randomly by dsDNA instability might be used as a new guide and direct DNAs cleavage.

The studies of the thermophilic Agos have revealed unique mechanistic features but still remain elusive in the feasibility of new gDNA in the canonical DNA cleavage reaction during the catalytic process. Of particular note biotechnologically, the nucleic acid-guided DNase activity of the CRISPR-associated protein has revolutionized the genome-detecting field recently [13-16], thus the thermophilic Agos were postulated suitable candidates for DNA reprograming applications [17, 18]. Here, we sought to address the mentioned question using *in vitro* assays with *Pf*Ago testing various gDNAs and substrates. To our considerable surprise, we observed that *Pf*Ago could stepwise cleave dsDNA substrates as guided by only a single gDNA, instead of one pair of gDNAs, in a precise and regulatory manner. After characterizing single gDNA directed stepwise dsDNA cleavage activity in detail, we developed a highly specific, multiplex detection platform which we used to distinguish four DNA targets in a convenient and efficient way, thereby demonstrating the utility of this Ago-based method for applications in biotechnology and molecular diagnostics.

We were initially interested in characterizing the cleavage performance of the hyperthermostable *Pf*Ago at 95 °C *in vitro*, in which the gDNAs were designed to target 600 bp PCR fragment as cleavage template sequences (Table S1, Fig. S1). Unexpectedly, we found that a single gDNA, rather than a pair of gDNAs, could mediate *Pf*Ago activity and rapidly produce cleavage products (Fig. 1A). When using short PCR fragment as the target templates (90 bp), *Pf*Ago was also able to cleave DNA in the presence of a single gDNA. It was surprisingly to observe distinct banding patterns for the cleavage products in reactions that used a single or a pair of gDNAs (Fig. 1B). These observations suggested that there may be some unusual enzymatic reaction process for this cleavage.

**Fig. 1.**
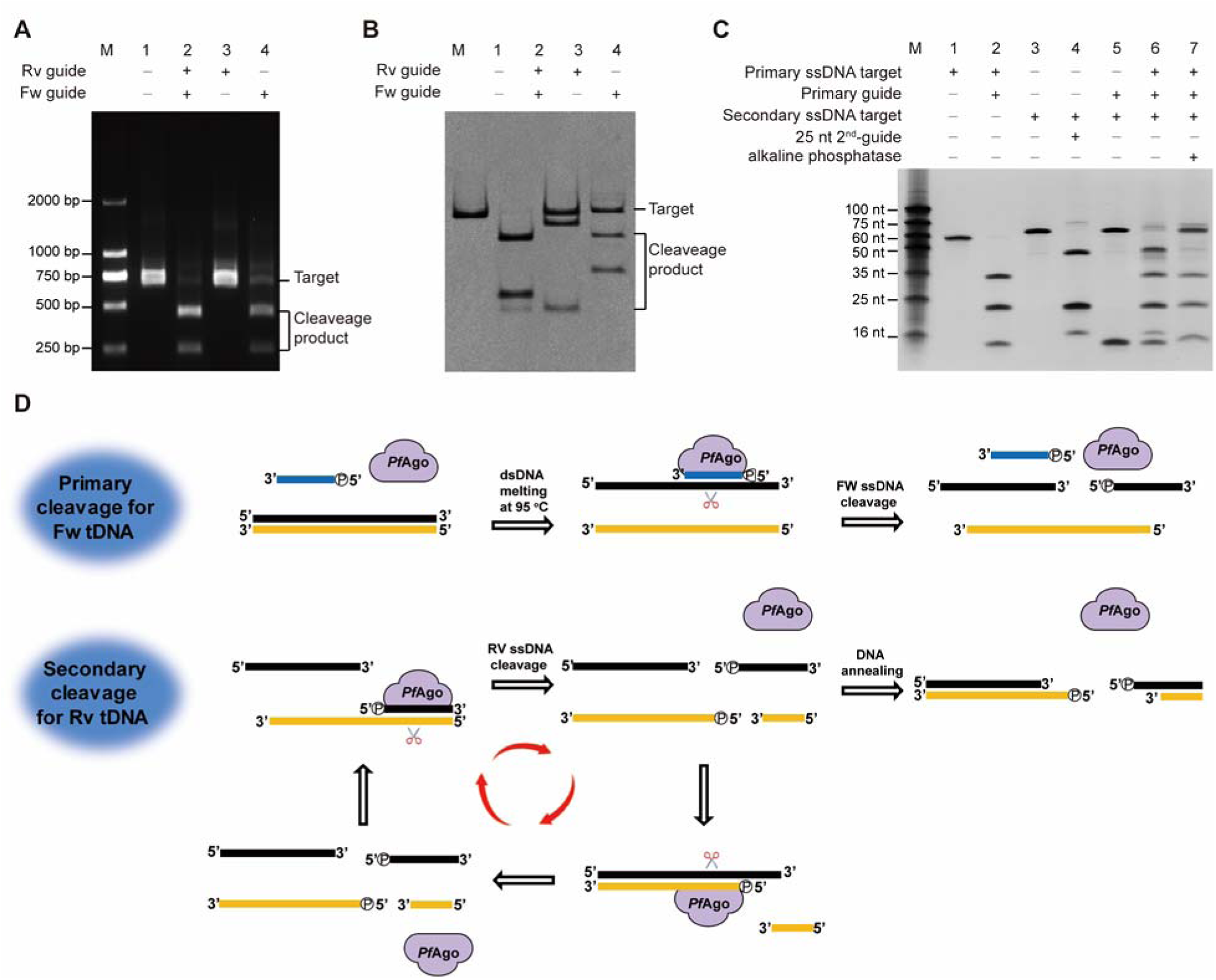
*Pf*Ago targeted DNA cleavage mediated by a single gDNA. (A) a single gDNA can mediate *Pf*Ago activity for 600 nucleotide (nt) DNA targets to produce DNA cleavage products. (B) a single gDNA can assist *Pf*Ago-mediated cleavage of 90 nt DNA targets, producing distinct banding patterns from reactions with a pair of gDNAs. (C) A cleavage fragment with a 5’-phosphate end can function as a renewed gDNA that can initialize an additional round of DNA cleavage if the reverse complement sequence of the original target is present. (D) Schematic of single gDNA-assisted *Pf*Ago-mediated activity for targeted DNA cleavage.

For such apparently off-target cleavage phenomenon, we proposed and first successfully confirmed that *Pf*Ago is able to bind the single gDNAs and form a guide-replacement process (Fig. S2). Then we conducted mass spectrometry analysis of products from *Pf*Ago-mediated reactions that used a single gDNA to cleave 60 nt ssDNA target. We detected two products: a 25 nt product with a 5’-phosphate end and a 35 nt product with a 3’-hydroxyl end (Fig. S3). Given that 5’-phosphate ends are known to be required for the activity of gDNAs in facilitating *Pf*Ago-mediated cleavage, our detection of cleaved ssDNA fragment with the 5’-phosphate ends raised the possibility that such fragments could potentially function as kind of “renewed-gDNA” for the further round of stepwise cleavage to the reverse complement sequences.

To probe the function of above cleavage products of the 60 nt ssDNA target, we added reverse complement 60 nt ssDNA to the reaction mixture from the primary reaction. Then we detected the expected 15 and 45 nt cleavage products as the cleavage of reverse complement ssDNA (Fig. 1C). Moreover, we found that enzymatic dephosphorylation treatment of the cleavage products from the primary reaction completely abolished any stepwise secondary cleavage activity for the reverse complement ssDNA. These results together support that cleavage fragments with 5’-phosphate ends generated in a primary cleavage can be used as the renewed-gDNA for stepwise rounds of cleavage targeting the reverse complement DNA sequence. Given that such subsequent rounds would also generate additional products with 5’-phosphate ends, we envisioned a ‘cascade’ of DNA cleavage rounds that targeted the two complementary sequences with each own of the corresponding new generated gDNAs (Fig. 1D).

We also investigated the kinetics of single gDNA-assisted *Pf*Ago-mediated cleavage using 30 nt ssDNA substrates labeled with fluorophore at their 5’-ends and fluorophore-quencher at their 3’-ends (FQ-ssDNA). The single gDNA-assisted *Pf*Ago-mediated cleavage activity, monitored by the fluorescence release quantitatively, fits the Michaelis-Menten equation with a catalytic efficiency (*k*_cat_/*K*_m_) of 7.1×10^7^ s^−1^M^−1^ (Fig. S4). To test the on-target cleavage specificity of target ssDNA, we designed multiple gDNAs targeting ssDNA and included mismatches across the length of the gDNA. These experiments showed that positions 9-12 of the gDNA sequence confer target specificity (Fig. 2A-B, Fig. S5), that is similar with mismatch tolerance pattern indicated by *Tt*Ago [10]. Moreover, these assays showed that di-nucleotide mismatches between gDNA and target sequences significantly decreased the cleavage activity of *Pf*Ago. These results suggested the feasibility of programing specificity of the cleavage to enable discrimination among target DNAs that differ by at least as little as di-nucleotide mismatches.

**Figure 2.**
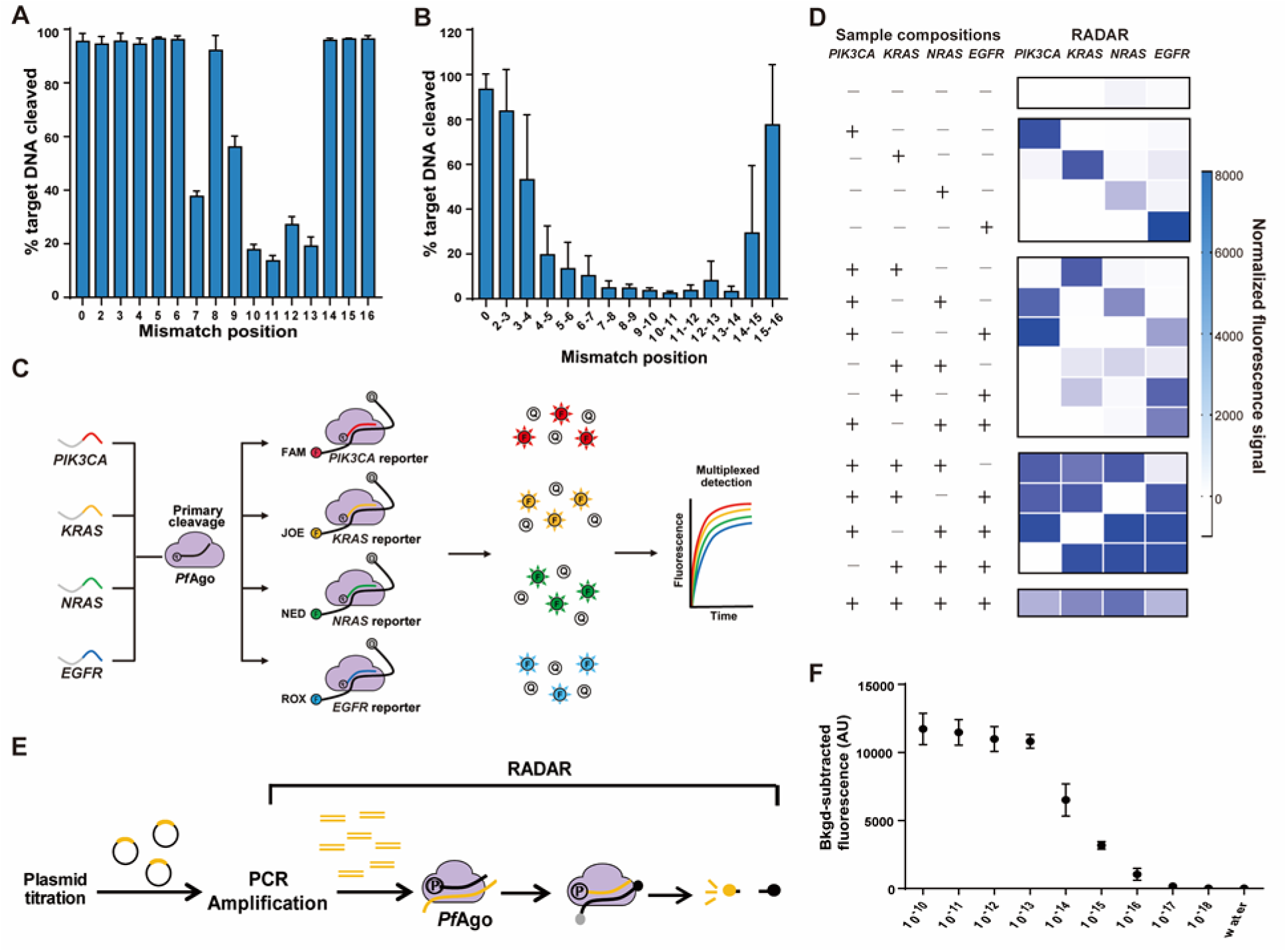
Single-gDNA-assisted *Pf*Ago-mediated DNA cleavage for highly specific and multiplexed nucleic acid detection. Observed activity for *Pf*Ago using a fluorophore labeled ssDNA with the indicated single nucleotide mismatches (A) and di-nucleotide mismatches (B). *Pf*Ago cleavage kinetics of ssDNA targets were measured, and the 20 min time point was plotted against the mismatch position. Ratios represent the average of three different targets measured in triplicate, and error bars represent mean ± s.d., where n = 9 (three replicates for three independent targets. (C) Schematic of one-pot four fluorophore multiplexing using orthogonal single-gDNA-assisted *Pf*Ago-mediated cleavage activity with each designed gDNA for respective ssDNA targets. (D) In-sample multiplexed detection of four ssDNA targets with corresponding designed gDNAs in an orthogonal manner.

Based on the single gDNA-assisted *Pf*Ago-mediated cleavage, we exploited it as a target DNA detection platform comprised the initial ssDNA templates, a designed gDNA targeting the template, *Pf*Ago, and reverse complement FQ-ssDNA reporter substrates. In this design, the gDNA will assist the *Pf*Ago-mediated cleavage of ssDNA template in the primary reaction thereby generate cleavage products to subsequently assist in the secondary cleavage of the FQ-ssDNA reporter substrates. We first systematic optimized the conditions in the single gDNA-assisted cleavage system, including gDNA length, the ratios of gDNA: *Pf*Ago: target DNA, along with the gDNAs tiling reporter length (Fig. S6-8). Afterall, we conducted the system based on single gDNA-assisted *Pf*Ago-mediated DNA cleavage to probe target ssDNA, in which set the gDNA (16 nt), ratios of *Pf*Ago: gDNA: target DNA (3:8:8), along with the gDNAs tiling reporter (30 nt).

We then extended experiments with a model system to test the ability to simultaneously discriminate amongst multiple different target sequences. These experiments tested four synthesized ssDNA templates (60 nt) based on the sequences of four different oncogenes (*PIK3CA, KRAS, NRAS*, and *EGFR*) (Fig. 2C). For fourplex detection, we used a PIK3CA-FAM reporter, a KRAS-JOE reporter, a NRAS-NED reporter, and an EGFR-ROX reporter, and we were able to detect each of the four targets in a single reaction (Fig. 2D, Fig. S9). Notably, no off-target signals were found in any of the reaction combinations, and the intensities of each of the four output signals in the multiplex reaction were at similar levels. We thus established proof-of-concept for the one-pot, multiplexed detection of multiple DNA substrates via orthogonal cleavage by *Pf*Ago.

The next application to expand the utility of such one-pot multiplexed detection would be adding PCR amplification for target sequences prior to subsequent *Pf*Ago-based detection. We tested this concept termed RADAR (<u>R</u>enewed gDNA <u>A</u>ssisted <u>D</u>NA cleavage with <u>Ar</u>gonaute; Fig. 2E), and evaluated the detection sensitivity as low as femtomolar level (Fig. 2E). Then we detected the presence of the four most common serotypes of the human papillomavirus (HPV): type 6, 11, 16, and 18 (Fig. 3A) [19]. First, we prepared samples containing various combinations of purified plasmids harboring serotype-specific alleles of the viral HPV *L1* gene (Fig. 3B). Then these samples were used as templates for amplification of approximately 110 bp amplicons using universal HPV primers. Subsequently, the amplified samples were reacted in a one-pot system containing four serotype-specific gDNAs and four serotype-specific FQ-ssDNA reporters. Establishing the combination of a PCR amplification step and our one-pot, multiplexed detection via orthogonal cleavage by *Pf*Ago, these RADAR experiments correctly and unambiguously identified each of the HPV subtypes present in each of the combination samples (Fig. 3C, S10).

**Figure 3.**
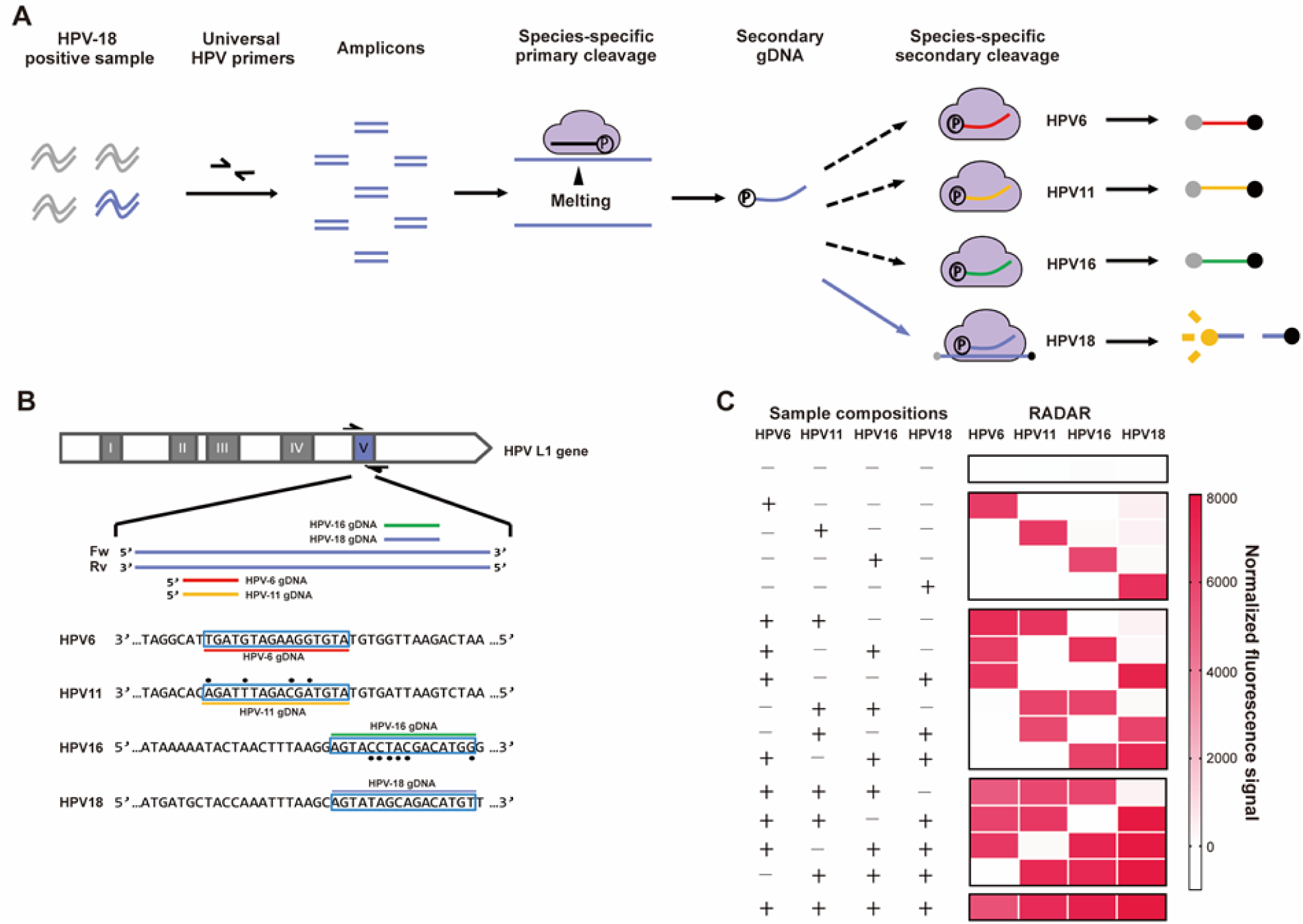
Multiplex RADAR and its application in HPV serotype detection. (A) Schematic of RADAR detection of one-pot four fluorophore multiplexing in an orthogonal manner. (B) The sequences for the four HPV serotypes within the hypervariable loop V of the *L1* gene targeted by gDNAs and *Pf*Ago; highlighted bases indicate the designed gDNA sequence. (C) RADAR detection of one-pot multiplexed detection of four dsDNA targets of HPV serotype with corresponding designed gDNAs.

Next, to apply RADAR for samples from human patients, we tested DNA samples extracted from human anal swabs that had been previously analyzed by a PCR-based method for HPV infection (Fig. S11). Within two hours, our RADAR method accurately identified HPV16 (9/9 agreement) and HPV18 (7/7 agreement) infections in the clinical samples with good correlation between the PCR-based intensity and RADAR signals (Fig. S11-12). We thusly demonstrate that our *Pf*Ago-cleavage based RADAR be used as a fast, sensitive, and reliable platform for the multiplexed detection of viral DNA in human samples.

In conclusion, our study first revealed a stepwise endonuclease activity of thermophilic *Pf*Ago which may provide a reprogrammable, efficient, and precise route for DNA editing. Our biochemical analysis has demonstrated that the single primary gDNA-directed dsDNA cleavage is the result of processive cleavage via precise generation of the kind of “renewed guide”. Offered by the stepwise cleavage and orthogonal target specificity of *Pf*Ago, we successfully established a one-pot multiplexed DNA detection platform as RADAR for availability in genotyping. Here, we anticipated RADAR as a versatile and sensitive method, especially for its advantage in cheap and accessible gDNA design, arbitrary cleaved target sequences, and operation simplicity in molecular diagnosis (Fig. S13). The exploration of thermophilic endonuclease make an alternative option for DNA editing thus would lead to a range of applications of medical diagnosis.

## Materials and methods

### Protein expression and purification

A synthesized codon-optimized version of the *Pf*Ago gene was ordered from GenScript (Nanjing, China), which was designed as the pET28a-derivative plasmid pEX-*Pf*Ago with an N-terminal His-tag. The expression plasmid was transformed into *E. coli* BL21(DE3) cells. A 5 mL seed culture was grown at 37 °C in LB medium with 50 μg/mL kanamycin, which was transferred to 1 L of LB in a shake flask containing 50 μg/mL kanamycin. The cultures were incubated at 37 °C until OD_600_ values of 0.8-1.0 was reached, and protein expression was then induced by addition of isopropyl β-D-thiogalactopyranoside (IPTG) to a final concentration of 1 mM, followed by incubation for 16 h at 20 °C. Cells were harvested for centrifugation for 20 min at 6,000 rpm, and the cell pellet was collected for later purification.

Cells pellet were resuspended in lysis buffer (20 mM Tris/HCl, 1 M NaCl, 2 mM MnCl2, pH 8.0), and then disrupted using a High Pressure Homogenizer at 600-800 bar for 3 min (Gefran, Italy). The lysates were centrifuged for 30 min at 12,000 rpm at 4 °C, after which the supernatants were used for Ni-NTA purification affinity with elution buffer (20 mM Tris/HCl, 1 M NaCl, 2 mM MnCl_2_, 200 mM imidazole, pH 8.0). Further gel filtration purification (Superdex 200, GE Tech) was carried out over a salt gradient from 250 mM to 2 M NaCl in elution buffer (20 mM Tris/HCl, 1 mM DTT, 5% glycerol, 2 mM MnCl_2_, pH 8.0). The resulting fractions from gel filtration were analyzed by SDS-PAGE, and fractions containing the protein were flashed frozen at −80 °C in storage buffer (20 mM Tris-HCl, pH 8.0, 250 mM NaCl, 0.5 mM MnCl_2_, 10% (v/v) glycerol).

### Nucleic acid preparation

The ssDNA target, gDNA, primers, and the FQ-reporters were synthesized commercially (Sangon Biotech, Shanghai, China). The 600 bp dsDNA target, were synthesized by GenScript (Nanjing, China) in the form of pET-28a derivative plasmids. The plasmid DNA was extracted using Plasmid DNA MiniPreps Kit (Generay, China). The 100 bp dsDNA target was obtained by amplification of the corresponding plasmids with designed primers using NCBI Primer-BLAST, with the option of amplicon size (between 90 and 120 nt), primer melting temperatures (between 54 °C and 67 °C), and primer size (between 20 and 25 nt). The PCR amplification used 2×PrimeSTAR Mix (Takara, Japan). The target dsDNA fragments for titration experiments were quantified using a PikoGreen dsDNA Quantitative Kit (Life iLab Biotech, China). All of the nucleic acids used in this study are listed in **Table S1**.

### DNA cleavage assays

Generally, *Pf*Ago-mediated cleavage assays were carried out in reaction buffer (15 mM Tris/HCl pH 8.0, 250 mM NaCl, and 0.5 mM MnCl_2_). For ssDNA cleavage, 0.2 μM *Pf*Ago, 2 μM gDNA, and 0.8 μM ssDNA target were mixed in reaction buffer and then incubated for 20 min at 95 °C in a thermocycler. Following high temperature incubation, the samples were cooled down by slowly lowering the temperature at the rate of 0.1 °C /s until it reached 10 °C. Reactions were stopped via the addition of loading Buffer (95% Formamide, 0.5 mmol/L EDTA, 0.025% bromophenol blue, 0.025% xylene cyanol FF) at a 1:1 ratio (v/v); the samples were then separated by 16% denaturing polyacrylamide gels and analyzed via staining with GelRed (Biotium, USA). The nucleic acids were visualized using a G:BOX Chemi imager (Syngene, USA) and quantitative analyzed by ImageQuant (GE Healthcare, USA).

For the 600 bp dsDNA cleavage assays, 0.16 μM *Pf*Ago, 2μM gDNAs, and 60 nM dsDNA target were mixed in reaction buffer; for the 95 bp dsDNA cleavage assays, 0.2 μM *Pf*Ago, 1 μM gDNAs, and 180 nM dsDNA target were mixed in reaction buffer before incubation for 15min at 95 °C in a thermocycler ((Eppendorf, Germany)). Following this high temperature incubation, the samples were cooled down by slowly lowering the temperature at a rate of 0.1 °C/s until it reached to 10 °C. Reactions were quenched with 5×DNA loading buffer (Generay, China) and analyzed by either 2% agarose gels or 15% non-denaturing polyacrylamide gels. Gels were stained and analyzed as described above.

For validation of 5’-phosphorylated end of cleavage fragment, totally 0.5 μM *Pf*Ago: 2 μM gDNAs: 2 μM primary ssDNA target were mixed in reaction buffer. Reactions were incubated for 35 min at 95 °C, then the cleavage products of primary ssDNA targets mediated by primary gDNAs were treated with or without alkaline phosphatase for further incubated with secondary ssDNA targets for 35min at 95 °C. Reactions were stopped by the addition of Loading Buffer (95% Formamide, 0.5 mmol/L EDTA, 0.025% Bromophenol Blue, 0.025% xylene cyanol FF) in a 1:1 ratio before the samples were resolved on 16% denaturing polyacrylamide gels. Gels were stained using GelRed (Biotium) and nucleic acids were visualized using a G:BOX Chemi imager (Syngene).

### Mass Spectrometry analysis for validation of the 5’-terminal end of cleavage fragments

For validation of the 5’-terminal end of cleavage fragment produced by gDNA-assisted *Pf*Ago activity, an *in vitro* cleavage reaction was set with ssDNA target cleaved by single gDNA. The *in vitro* assay consisted of 0.5 μM *Pf*Ago, 2 μM single gDNA, and 2 μM of the ssDNA target in a total volume of 100 µL; for control reactions, single gDNA was replaced by UltraPure water. The cleavage reactions were incubated at 95 °C for 1 h, and reaction products were purified using an ethanol precipitation method in which a one-tenth volume of 3 M NaOAc and three volumes of ice cold ethanol were added; the samples were kept at −20 °C for 2 hours, and were subsequently centrifuged for 15 minutes at 12,000 rpm. The supernatant was removed, and two volumes of 80% ethanol were then added and incubated −20°C for 2 hours. The supernatant was decanted, and samples were centrifuged for 5 minutes at 12,000 rpm. After air-drying the pellet, samples were resuspended in 50 μL of UltraPure water.

A Model 1290 Ultra Performance LC (UPLC) system (Agilent Technologies) with a Zorbax XDB C8 reverse phase column (4.6×150 mm, 5 μm, particle size, Agilent Technologies) was used for sample separate, with the column temperature controlled at 30 °C. The two mobile phase solvents consisted of buffer A as H_2_O with 0.1% ammonium hydroxide (v/v) and buffer B as acetonitrile. The flow rate of the mobile phase was 0.4 ml/min, and the injection volume was 5μl. LC flow was coupled to an Agilent model 6230 accurate mass time-of-flight (TOF) MS (Agilent Technologies) equipped with an Agilent Jet Stream electrospray ionization (ESI) source. The mass spectrometer was operated in negative ion mode with the following parameters: capillary voltage, 3500 V; the skimmer voltage, 65 V; nozzle voltage, 800 V; fragmentor, 135 V. Nitrogen was used as the drying (8 L/min, 325 °C), sheath (11 L/min, 350 °C), and nebulizer gas(35 psi). The mass spectrometer was tuned for large MW ions, and data was acquired between m/z 400-5000. Data was saved in centroid mode, using Agilent MassHunter Workstation Data Acquisition Software (revision B.04); the MaxEnt deconvolution algorithm was used to generate a calculated neutral mass spectrum from the negatively charged ion data.

### EMSA assays

For the EMSA assays, 1 μM *Pf*Ago was pre-incubated with 0.2 μM fluorescent labeled gDNAs for 5 min at 95 °C in reaction buffer (15 mM Tris/HCl pH 8.0, 250 mM NaCl, and 0.5 μM MnCl_2_,). The subsequent gDNAs without fluorescent labelling were then added at various concentrations (0, 0.2, 0.4, 0.8, 2 μM). The reaction mixtures were incubated for another 5 min. Next, gel loading buffer (2.5% Ficoll 400, 62.5 mM Tris/HCl, pH 6.8) was added to quench the reaction, and the samples were analyzed by 8% non-denaturing polyacrylamide gels. Subsequently, the fluorescent labeled gDNA and the protein–gDNA complexes were visualized on a Fuji FLA7000 scanner with fluorescence measurements of λ_ex_ at 535 nm and λ_em_ at 595 nm.

### Fluorophore quencher (FQ)-labeled reporter assays

The *Pf*Ago cleavage assays with FQ-labeled ssDNA were carried out in reaction buffer (15 mM Tris-HCl, 250 mM NaCl, 0.5 mM MnCl_2_, pH 8.0). For the FQ-labeled ssDNA as indicator of primary cleavage, the reactions were mixed to final concentration of 300 nM *Pf*Ago, 2 µM gDNA, with 600 nM ssDNA-FQ reporter substrates in a 25 µL total reaction volume. For measuring the effect of gDNA concentration on secondary cleavage efficiency, the reactions were mixed to final concentration of 300 nM *Pf*Ago, 800 nM target ssDNA, and 600 nM ssDNA-FQ reporter substrates in a 25 µL total reaction volume; the gDNA was added separately to final concentrations of 0 µM, 0.4 µM, 0.8 µM, 1.2 µM, 1.6 µM, 2 µM, 4 µM, and 8 µM. For measuring the effect of target ssDNA concentration on secondary cleavage efficiency, the reactions were mixed to final concentration of 300 nM *Pf*Ago, 800 nM gDNA, and 600 nM ssDNA-FQ reporter; target ssDNA was added separately to final concentrations of 0 nM, 40 nM, 80 nM, 200 nM, 320 nM, 400 nM, 600 nM, and 800 nM. For measuring the effect of the *Pf*Ago concentration on secondary cleavage efficiency, the reactions were mixed to final concentration of 800 nM target ssDNA, 800 nM gDNA, and 600 nM ssDNA-FQ reporter; *Pf*Ago was added separately to final concentrations of 0 nM, 40 nM, 80 nM, 200 nM, 320 nM, 400 nM, 600 nM, and 800 nM. Reactions (25 µL, Axygen qPCR tube) were incubated in the Mastercycler® real-plex instrument (Eppendorf, Germany) for up to 30 minutes at 95 °C.

For multiplex detection of ssDNA targets, reactions were performed as follows: the components of the reaction were 1.2 µM *Pf*Ago: 0.4 µM gDNA for each DNA: 0.4 µM target ssDNA for each DNA, and the ssDNA-FQ reporter was added in the following concentrations: 0.4 µM FAM-labeled ssDNA-FQ reporter; 0.8 µM JOE-labeled ssDNA-FQ reporter; 1.2 µM NED-labeled ssDNA-FQ reporter; 2 µM ROX-labeled ssDNA-FQ reporter. Reactions (25 µL, Axygen qPCR tube) were incubated in the Mastercycler® real-plex instrument (Eppendorf, Germany) for up to 30 minutes at 95 °C, with fluorescence measurements taken every 30 seconds (FAM-labeled ssDNA-FQ reporter λ_ex_: 495 nm; λ_em_: 520 nm, JOE-labeled ssDNA-FQ reporter λ_ex_: 529 nm; λ_em_: 550 nm, NED-labeled ssDNA-FQ reporter λ_ex_: 557 nm; λ_em_: 580 nm, ROX-labeled ssDNA-FQ reporter λ_ex_: 586 nm; λ_em_: 605 nm).

For the dsDNA secondary cleavage analysis, the dsDNAs were prepared from HPV-16 containing plasmids by PCR. Briefly, 25 µL reactions containing 0.5 µL plasmids (290 ng/ µL), 0.6 µM forward and reverse primer, 2×AceQ qPCR Probe Master Mix (Vazyme Biotech, China) were amplified for target dsDNA fragment. The dsDNA pruduct (2 pM) was mixed with 0.4 µM FAM-labeled ssDNA-FQ reporter, 1.2 µM *Pf*Ago, 0.5 mM MnCl_2_ and 2 µM reverse gDNA (targeting the secondary gDNA production chain) or both reverse gDNA and forward gDNA. Reactions were incubated in the Mastercycler® real-plex instrument (Eppendorf, Germany) for up to 30 minutes at 95 °C with fluorescence measurements taken every 30 seconds (ssDNA-FQ reporter λ_ex_: 495 nm; λ_em_: 520 nm).

For the Michaelis-Menten analysis, a total of 0.8 μM *Pf*Ago, 0.8 μM gDNA was mixed in reaction buffer, and the reactions were initiated by adding 0.04, 0.08, 0.2, 0.4, 0.6, or 0.8 µM of the substrate (FQ-labeled ssDNA); reactions were incubated in the Mastercycler® real-plex instrument (Eppendorf, Germany) for up to 990 seconds at 95 °C with fluorescence measurements taken every 10 seconds (λ_ex_: 535 nm; λ_em_: 595 nm).

### Multiple ssDNA detection

The 4 channel multiplex ssDNA detection system was established for 4 different types of target gene detection in a one-pot reaction system. The reaction system was similar with the aforementioned FQ-labeled reporter assays, with the following modifications: 25 µL reactions contained 400 nM target ssDNA (*PIK3CA, KRAS, NRAS*, and *EGFR*), 0.4 µM gDNAs (ssDNA1, ssDNA2, ssDNA3, ssDNA4 targeting), 2.4 µM *Pf*Ago, 0.4 µM FAM-labeled ssDNA-FQ reporter (ssDNA1), 0.8 µM JOE-labeled ssDNA-FQ reporter (ssDNA2), 1.2 µM NED-labeled ssDNA-FQ reporter (ssDNA3), and 2 µM ROX-labeled ssDNA-FQ reporter (ssDNA4). Reactions were incubated in the Mastercycler® real-plex instrument (Eppendorf, Germany) for up to 30 minutes at 95 °C with fluorescence measurements taken every 30 seconds (FAM-labeled ssDNA-FQ reporter λ_ex_: 495 nm; λ_em_: 520 nm, JOE-labeled ssDNA-FQ reporter λ_ex_: 529 nm; λ_em_: 550 nm, NED-labeled ssDNA-FQ reporter λ_ex_: 557 nm; λ_em_: 580 nm, ROX-labeled ssDNA-FQ reporter λ_ex_: 586 nm; λ_em_: 605 nm).

### RADAR assays

HPV detection assays were performed as above, with the following modifications: 50 µL reactions contained 0.5 µL plasmids of each gene (40 ng/µL), 0.5 µM universal primers. 0.4 µM HPV-6, HPV-11, HPV-16, HPV-18 targeting gDNAs, 2.4 µM *Pf*Ago, 0.4 µM FAM-labeled ssDNA-FQ reporter (HPV-16), 0.8 µM JOE-labeled ssDNA-FQ reporter (HPV-18), 1.2 µM NED-labeled ssDNA-FQ reporter (HPV-6), and 2 µM ROX-labeled ssDNA-FQ reporter (HPV-11) were added into the PCR reaction mixtures. Reactions (35 µL final volume) were incubated in Reactions (25 µL, Axygen qPCR tube) were incubated in the Mastercycler® real-plex instrument (Eppendorf, Germany). Initially, 30 cycles of PCR amplification were conducted (95 °C, 30 s; 60 °C, 20 s), then incubated for 30 minutes at 95 °C with fluorescence measurements taken every 30 seconds (FAM-labeled ssDNA-FQ reporter λ_ex_: 495 nm; λ_em_: 520 nm, JOE-labeled ssDNA-FQ reporter λ_ex_: 529 nm; λ_em_: 550 nm, NED-labeled ssDNA-FQ reporter λ_ex_: 557 nm; λ_em_: 580 nm, ROX-labeled ssDNA-FQ reporter λ_ex_: 586 nm; λ_em_: 605 nm).

For evaluated the detection sensitivity of RADAR, the template of the HPV-16 plasmid was diluted to 10^−10^ M, 10^−11^ M, 10^−12^ M, 10^−13^ M, 10^−14^ M, 10^−15^ M, 10^−16^ M, 10^−17^ M, 10^−18^ M, 10^−19^ M, or 10^−20^ M. With the same detection system, the templates of this concentration gradient were analyzed separately, with the detection values normalized to the maximum mean fluorescence signal. For the HPV clinical sample identification by RADAR, the detection method was the same as above, with the detection of HPV types 16 or 18 in human samples implemented using *Pf*Ago targeting the hypervariable loop V of the L1 gene within HPV-16 or HPV-18. A one-way ANOVA with Dunnett’s post-test was used to determine the positive cutoff (set at p ≤ 0.05) for the identification of HPV-16 or HPV-18 in patient samples.

### Human clinical sample collection, DNA preparation and validation

Donors for providing anal swab samples were recruited from the Fifth People’s Hospital of Shanghai, Fudan University (Shanghai). The study was approved by Human Research Committee of the Fifth People’s Hospital of Shanghai, Fudan University (Shanghai). The samples were collected from an anal swab into a Thinprep™ vial with 1 ml sterilizing saline for HPV testing. The cell suspension was centrifuged for 5 minutes at 12,000 rpm and cell precipitate was resuspend in 1 ml sterilizing saline used for DNA extraction with TIANamp Genomic DNA kit (TianGen Biotechnologies, Beijing). Subsequently, 5 μL of DNA was used for the HPV consensus PCR analysis and the RADAR test in parallel.

The PCR-based HPV genotyping and validation was performed using a commercially-available kit of Liferiver HPV Genotyping Real Time PCR kit (ZJ Bio-Tech Corporation, Shanghai). The principle of this method is based on the sequence specific probe for taqman real time detection. After 40 amplification cycles, specimens were probed with a FAM-labeled specific probe mixture. Specimens negative for beta-globin gene amplification were excluded from analysis. The results were recorded based on the Ct value (<38) and amplification curve (S-shape) of the real time PCR. Samples with results recorded as 1 or more were considered to be positive.

## Acknowledgments

We thank Professor Yanli Wang from institute of Biophysics, Chinese Academy of Sciences for comments and discussions. This research was supported in part by the grants from Natural Science Foundation of China (31770078) and Ministry of Science and Technology (2017YFE0103300).

## Conflict of Interest

Shanghai Jiao Tong University has applied for a patent (China application no. 201810291873.0) on RADAR with Y.F., G.X., Q.L., and Y.C. listed as co-inventors.

## Supplementary Materials

Figures S1-S12

Tables S1-S7

