## Supplemental Material for "The stepwise endonuclease activity of a thermophilic Argonaute protein"

**Affiliations:**

**Content**

**Supplementary Figures:**

Figure S1. Purification of *Pf*Ago protein.

Figure S2. Detection dynamic process of *Pf*Ago binding gDNA using an electrophoresis mobility shift assay (EMSA).

Figure S3. TOF-MS analysis of cleavage ends from reaction of *Pf*Ago with ssDNA target.

Figure S4. Michaelis-Menten analysis reveals single gDNA-assisted *Pf*Ago-mediated cleavage activity with a ssDNA-FQ reporter.

Figure S5. The proximal mismatches of gDNA with dsDNA target in the assay of fluorescent reporter provide specificity for single gDNA-assisted *Pf*Ago-mediated cleavage activity.

Figure S6. Cleavage efficiency influenced by gDNA length.

Figure S7. Efficiency and specificity of single gDNA-assisted *Pf*Ago-mediated cleavage activity.

Figure S8. Testing single gDNA-assisted *Pf*Ago-mediated cleavage activity effected by gDNAs tiling reporter.

Figure S9. Multiplex detection based on orthogonal single gDNA-assisted *Pf*Ago-mediated cleavage activity with ssDNA targets.

Figure S10. Multiplex detection of 4 subtypes HPV containing plasmids by RADAR. Figure S11. Identification of HPV serotypes by PCR based (left) and RADAR detection (right).

Figure S12. RADAR analysis of HPV types 16 and 18 in patient samples.

**Supplementary Tables:**

Table S1: *Pf*Ago protein used in this study.

Table S2. gDNA used in this study.

Table S3. ssDNA targets used in this study.

Table S4. primers used in this study.

Table S5. Cleavage reporters used in this study.

Table S6. Plasmids used in this study.

Table S7. Nucleic acid for EMAS used in this study.

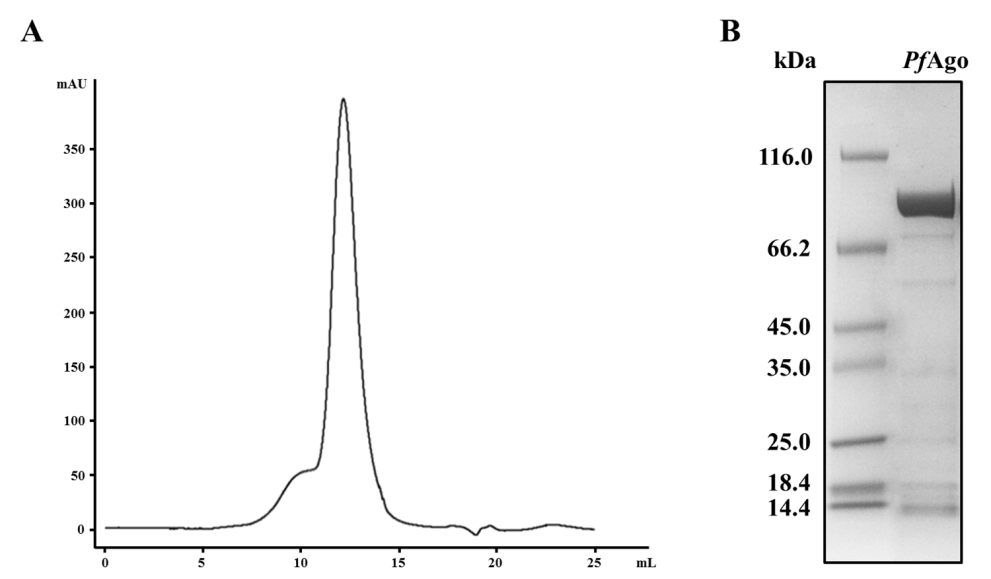

**Figure S1. Purification of *Pf*Ago protein.** (A) Chromatograms of size exclusion chromatography for *Pf*Ago. Measured UV absorbance (mAU) at 280 nm is shown against the elution volume (ml). (B) Final SDS-PAGE gel of *Pf*Ago protein used in this study. The calculated molecular weight of *Pf*Ago is 90.4 kDa.

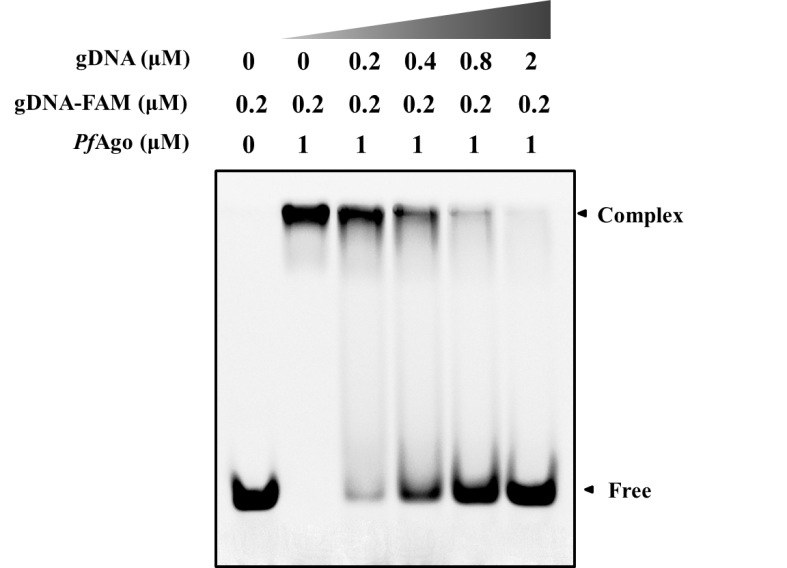

**Figure S2. Detection dynamic process of *Pf*Ago binding gDNA using an electrophoresis mobility shift assay (EMSA)**. Initially, *Pf*Ago was incubated with fluorophore labeled gDNA, and then various concentrations (0, 0.2, 0.4, 0.8 and 2 µM) of non-labeled gDNA was harnessed in the reaction to examine the binding affinities of *Pf*Ago with previous labeled gDNA.

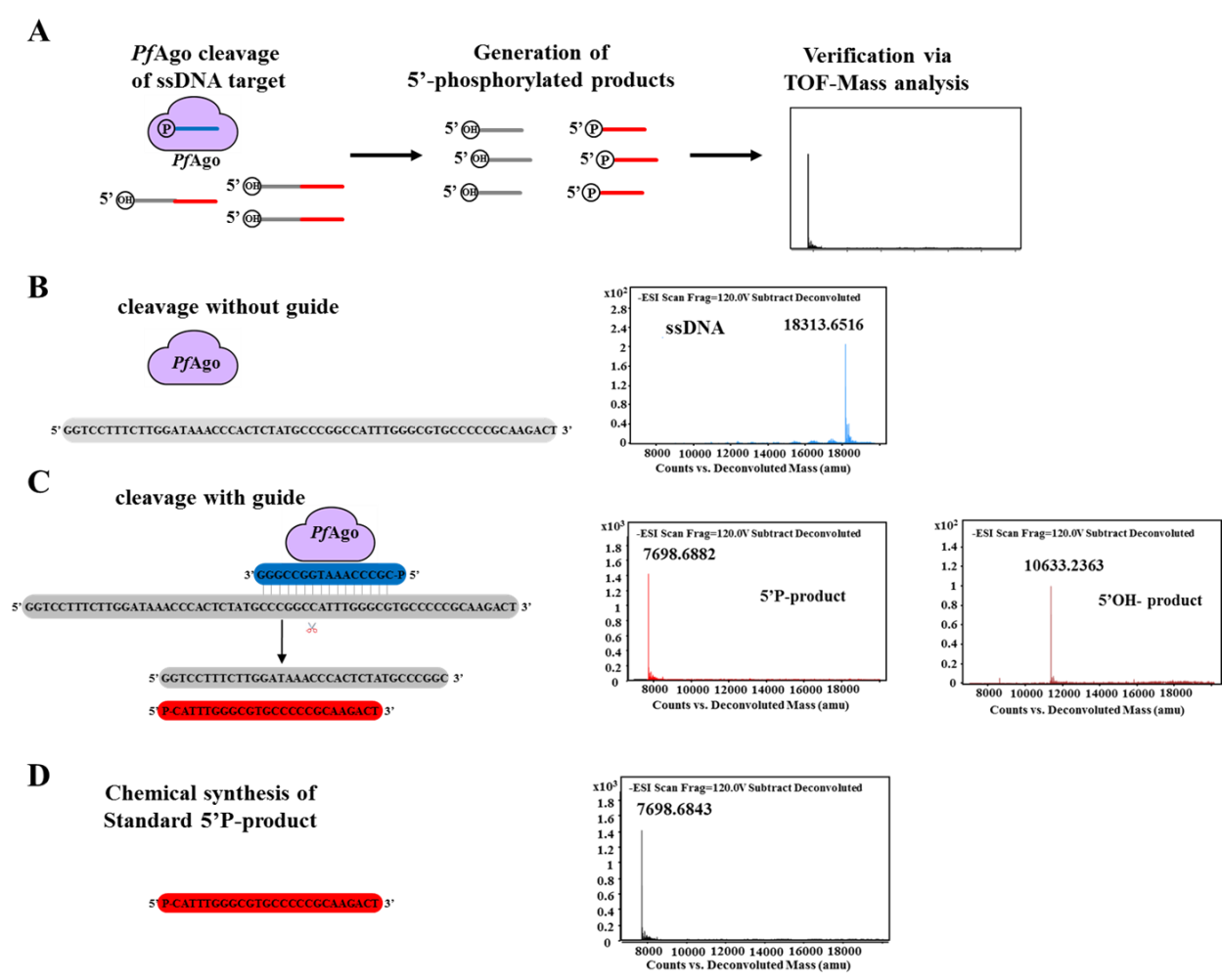

**Figure S3. TOF-MS analysis of cleavage ends from reaction of *Pf*Ago with ssDNA target.** A) Schematic of *Pf*Ago specific cleavage with ssDNA target and mass spectrometric analysis to verify cleavage products. Mass spectrometry analysis of cleavage products from *Pf*Ago in vitro assay without guide (B) and with guide (C). (D) Chemical synthesis of cleavage product used as the standard. The ESI-MS signals were measured in negative single ion mode, and the deconvolution mass are labeled.

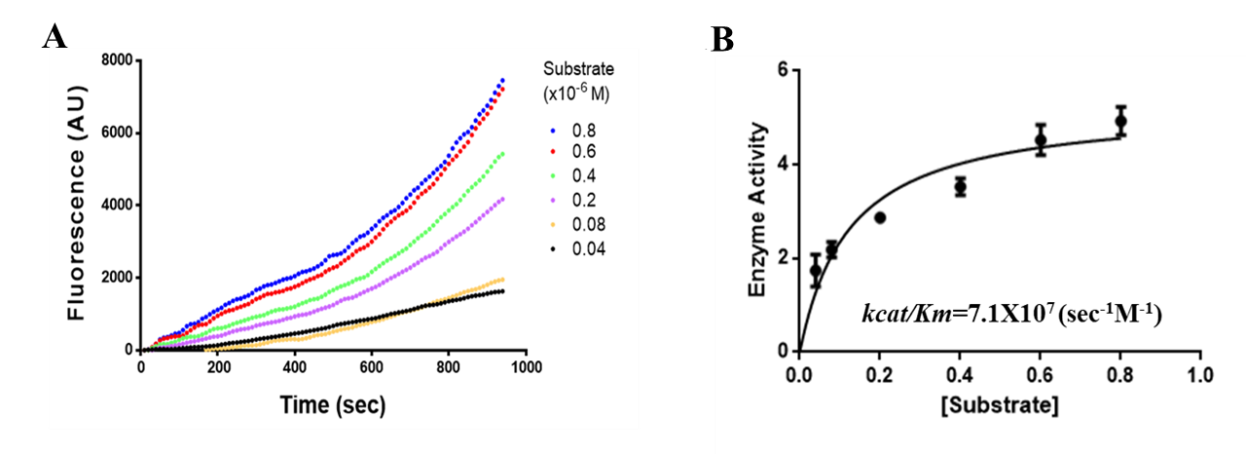

**Figure S4. Michaelis-Menten analysis reveals single gDNA-assisted *Pf*Ago-mediated cleavage activity with a ssDNA-FQ reporter.** (A) Representative plots of initial velocity versus time for a ssDNA reporter, using 0.6 nM *Pf*Ago and increasing fluorophore labeled ssDNA substrate concentrations at 95°C. (B) Michaelis-Menten fits for the ssDNA reporter. Calculated *k_cat_*, *K_m_* and *k_cat_/K_m_* values are expressed as the mean ± s.d., where n = 3 replicates.

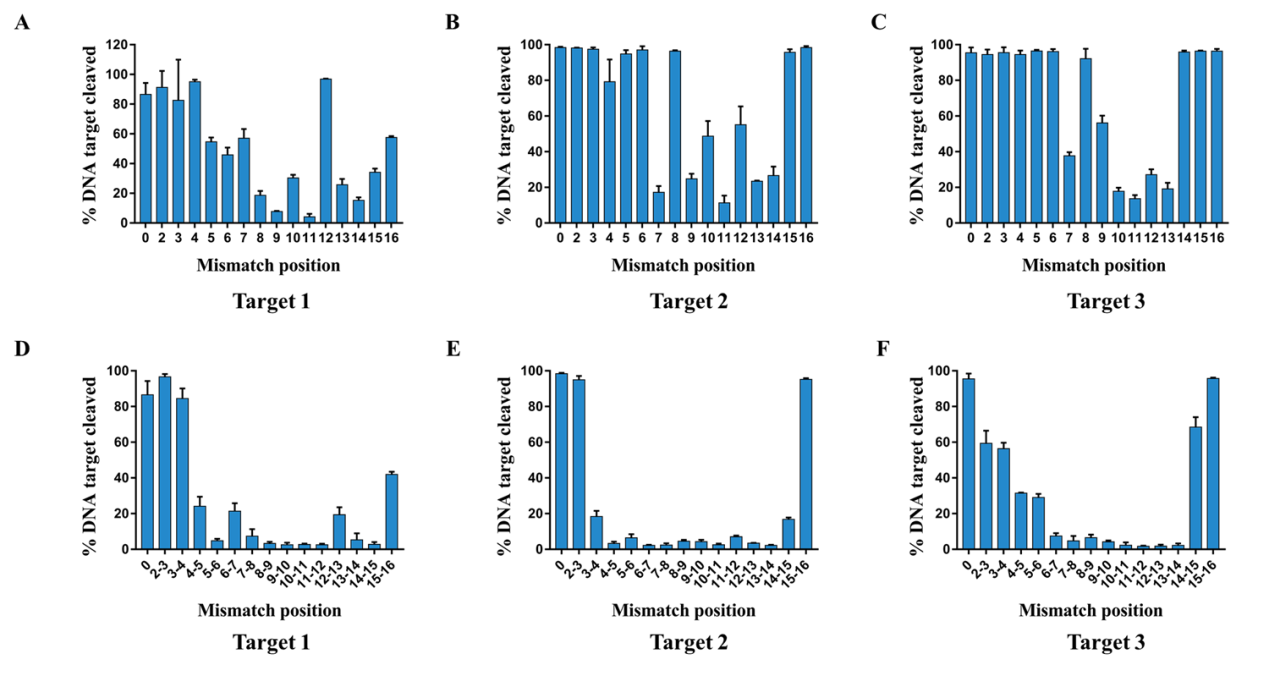

**Figure S5. The proximal mismatches of gDNA with dsDNA target in the assay of fluorescent reporter provide specificity for single gDNA-assisted *Pf*Ago-mediated cleavage activity.** (A) to (C) Quantification of cleavage kinetics using mismatched gDNAs with single nucleotide for three distinct target sequences of JHF1, KRAS, PIK3CA, respectively; (D) to (F) Quantification of cleavage kinetics using mismatched gDNAs with continuous two nucleotide for three distinct target sequences of JHF1, KRAS, PIK3CA, respectively. Error bars represent the mean ± s.d., where n = 3 replicates.

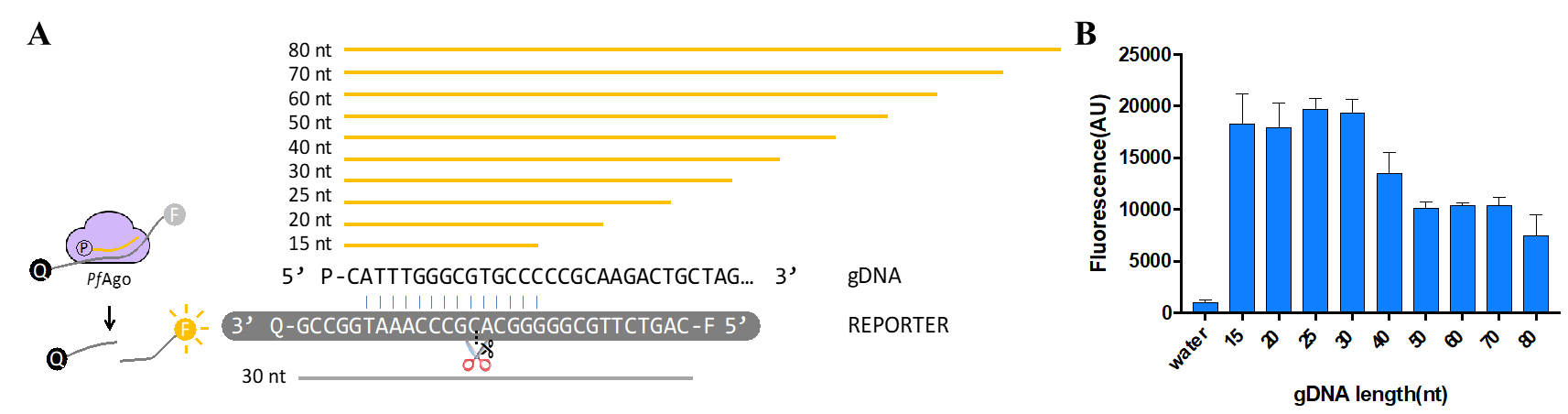

**Figure S6. Cleavage efficiency influenced by gDNA length.** (A) Schematic of pairing overlap between gDNA and fluorophore labeled ssDNA target. (B) Fluorophore labeled ssDNAs were catalyzed by different length gDNA-mediated-*Pf*Ago. Each bar represents the mean fluorescence signal from fluorophore labeled ssDNA cleavage after 40 min at 95℃. Error bars represent the mean ± s.d., where n = 3 replicates.

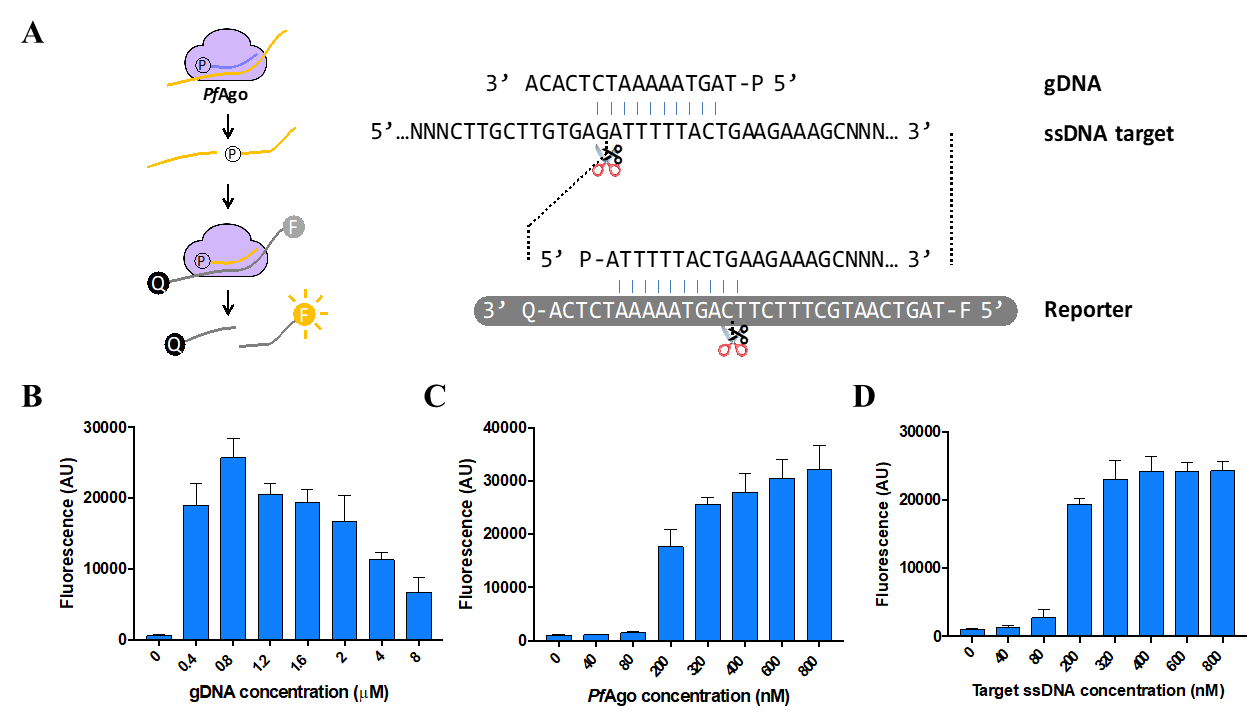

**Figure S7. Efficiency and specificity of single gDNA-assisted *Pf*Ago-mediated cleavage activity.** (A) Schematic of assay determined single gDNA-assisted *Pf*Ago-mediated cleavage activity. (B) single gDNA-assisted *Pf*Ago-mediated cleavage activity with gDNA of varying concentration when *Pf*Ago at 0.3 µM and target ssDNA at 0.8 µM. (C) single gDNA-assisted *Pf*Ago-mediated cleavage activity of *Pf*Ago with ssDNA of varying concentration when *Pf*Ago at 0.3 µM and gDNA at 0.8 µM. (D) single gDNA-assisted *Pf*Ago-mediated cleavage activity of *Pf*Ago with *Pf*Ago of varying concentration when gDNA at 0.8 µM and target ssDNA at 0.8 µM. Each bar represents the mean fluorescence signal from fluorophore labeled ssDNA cleavage after 40 min at 95℃. Error bars represent the mean ± s.d., where n = 3 replicates.

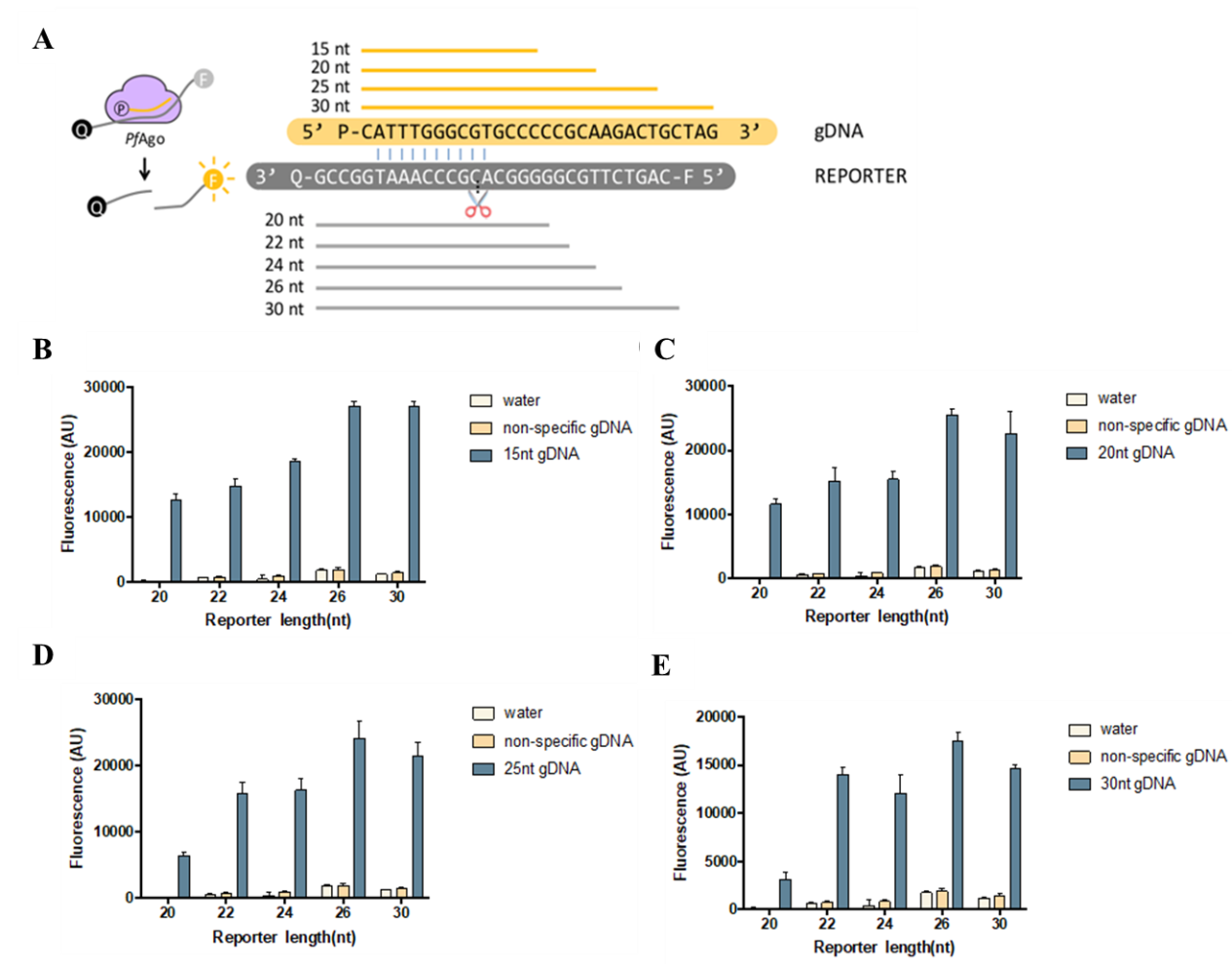

**Figure S8. Testing single gDNA-assisted *Pf*Ago-mediated cleavage activity effected by gDNAs tiling reporter.** (A) Schematic of locations tiled gDNA targeting reporter. (B) Single gDNA-assisted *Pf*Ago-mediated cleavage activity with 15 nt gDNA tiled across reporters of varying lengths. (C) Single gDNA-assisted *Pf*Ago-mediated cleavage activity with 20 nt gDNA tiled across reporters of varying lengths. (D) Single gDNA-assisted *Pf*Ago-mediated cleavage activity with 25 nt gDNA tiled across reporters of varying lengths. (E) Single gDNA-assisted *Pf*Ago-mediated cleavage activity with 15 nt gDNA tiled across reporters of varying lengths. Error bars represent the mean ± s.d., where n = 3 replicates.

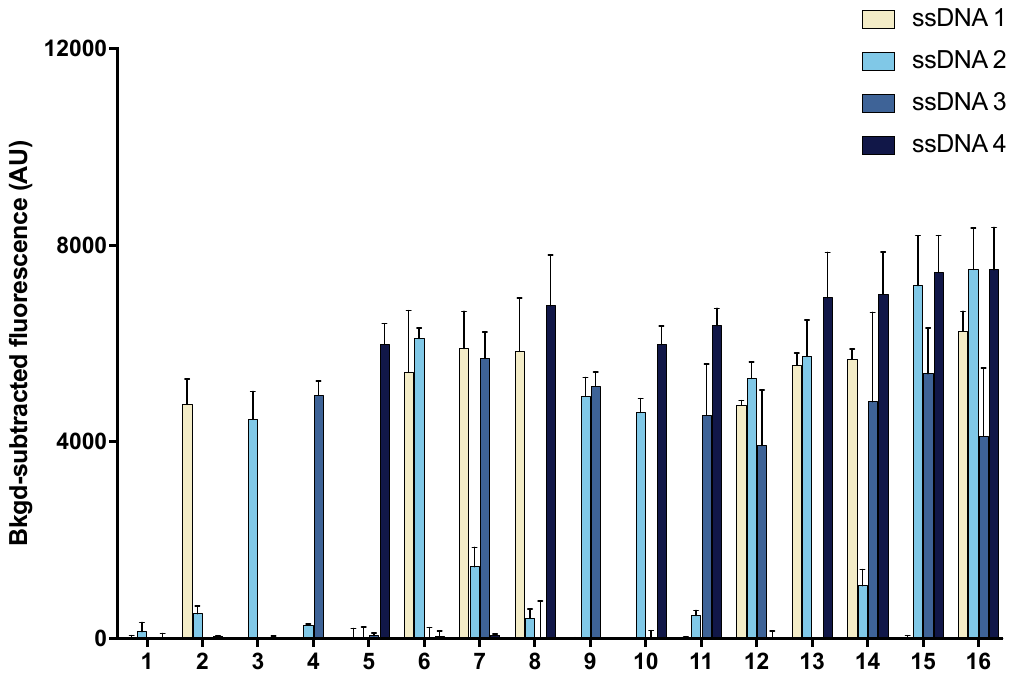

**Figure S9. Multiplex detection based on orthogonal single gDNA-assisted *Pf*Ago-mediated cleavage activity with ssDNA targets**. Detection of four ssDNA targets by orthogonal single gDNA-assisted *Pf*Ago-mediated cleavage activity in a 16 combination samples; input of water was used as a control for background deduction. Error bars represent mean ± s.d., where n = 3 replicates.

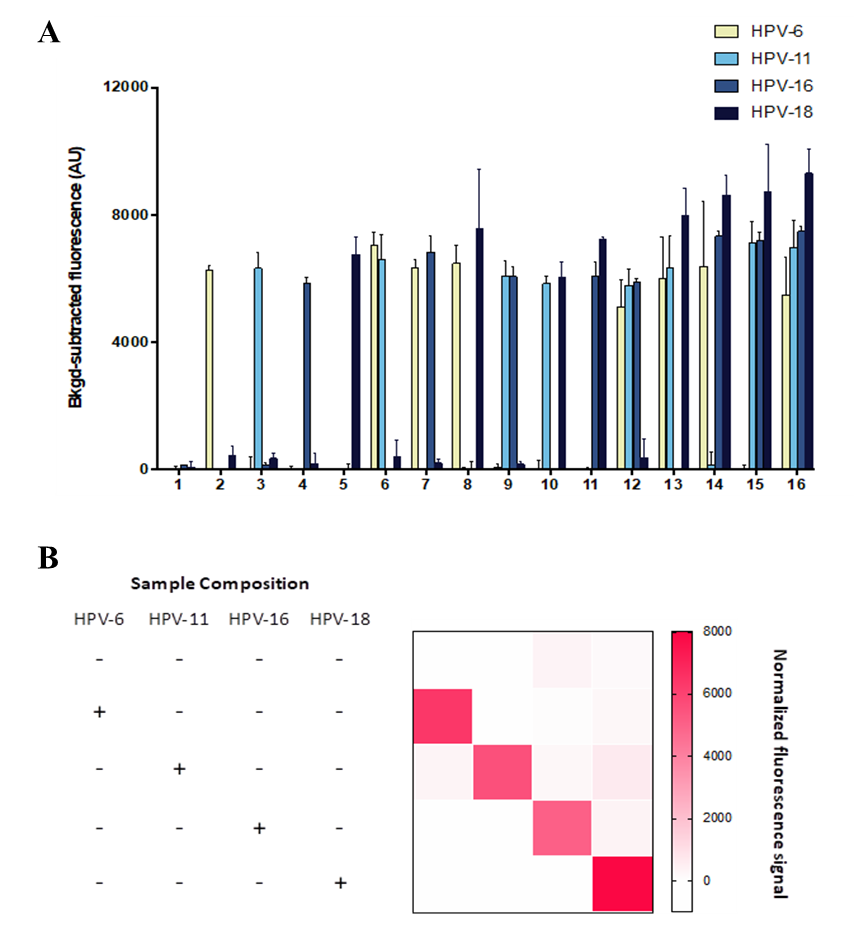

**Figure S10. Multiplex detection of 4 subtypes HPV containing plasmids by RADAR.** (A) Collateral RADAR system on four types of dsDNA input in 16 combination samples; input of water (no ssDNA) was used as a control. The absence of fluorescence signal in specimens that were not detected with all of those four subtypes ssDNA, was an indicator of good specificity by RADAR. Error bars represent mean ± s.d., where n = 3 replicates. (B) Heatmap depiction of RADAR results.

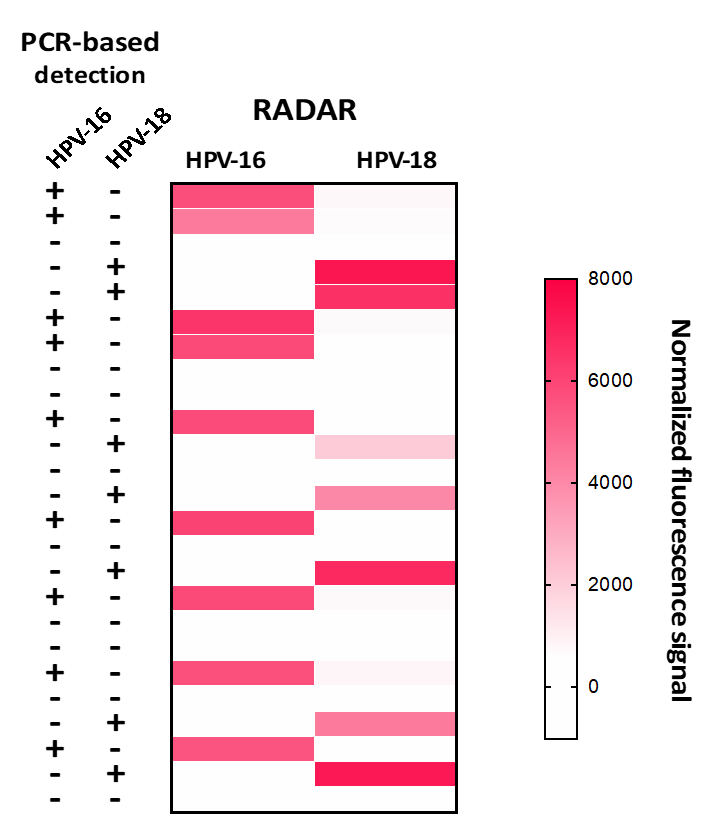

**Figure S11. Comparion of HPV serotypes identification by PCR based (left) and RADAR detection (right)**. The RADAR heatmap represents the normalized mean fluorescence values.

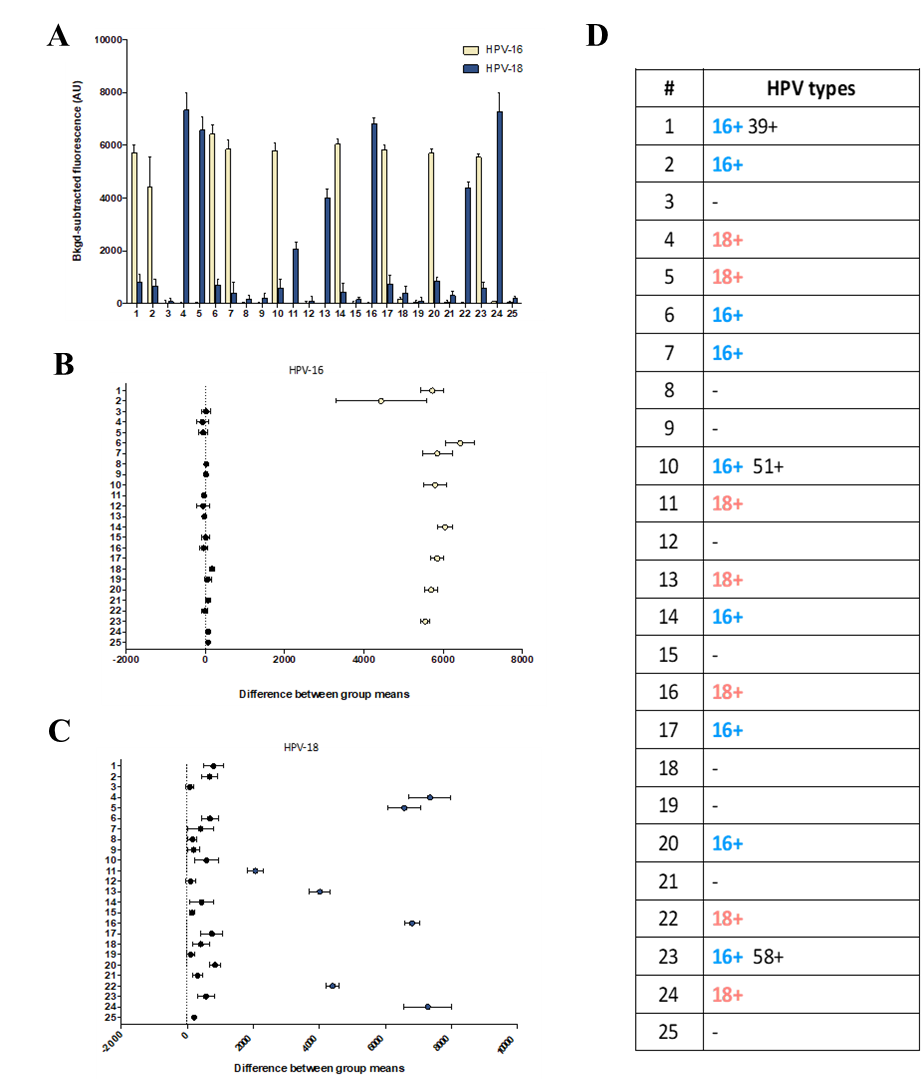

**Figure S12. RADAR analysis of HPV types 16 and 18 in patient samples** (A) Detection of HPV types 16 or 18 by RADAR in human anal clinical samples; input of water (no HPV) was used as a control. The absence of fluorescence signal in specimens that were not infected with HPV types 16 or 18, but did contain other HPV types, was an indicator of good specificity by RADAR. Error bars represent mean ± s.d., where n = 3 replicates. Plot of 95% confidence intervals of difference between control and sample groups of HPV types 16 (B) and 18 (C), based on a one-way ANOVA with Dunnett’s post test, where n = 3 replicates. Highlighted sample numbers indicate positive detection of HPV16 (left) or HPV18 (right) in patient samples, where **p≤0.01 and ***p≤0.001. (D) Summary of PCR-based detection of HPV types 16 (column 2 and yellow circles) and 18 (column 3 and orange circles) and identification of other HPV types by PCR in patient samples (column 4); subjective intensive values (0–4 scale) were assigned for each PCR based validation (columns 2 and 3).

**Table S1. *Pf*Ago protein purified in this study.**

| Protein name | Strain name | Accession number |
| --- | --- | --- |
| *Pf*Ago | *Pyrococcus furiosus* | WP_011011654.1 |

**Table S2. gDNA used in this study.**

| Name | Complete gDNA sequence | Target | Presence |
| --- | --- | --- | --- |
| JHF-RV-gDNA | P-TGCCCAAATGGCCGGG | JHF1-600bp dsDNA | Fig.1A |
| JHF-FW-gDNA | P-TATGCCCGGCCATTTG | JHF1-600bp dsDNA | Fig.1A |
| PIK3CA-RV-gDNA | P-TTCTCCTGCTCAGTGA | PIK3CA-95bp dsDNA | Fig.1B |
| PIK3CA-FW-gDNA | P-TGAAATCACTGAGCAG | PIK3CA-95bp dsDNA | Fig.1B |
| Primary gDNA | P-TGCCCAAATGGCCGGG | Primary ssDNA | Fig.1C/Fig.S3C |
| 25nt 2^nd^ gDNA | P-CATTTGGGCGTGCCCCCGCAAGACT | Secondary ssDNA | Fig.1C/Fig.S3D |
| Target 1_gDNA_2 | P-TCCCCAAATGGCCGGG | ssDNA 1 | Fig.2A/Fig.S5A |
| Target 1_gDNA_3 | P-TGGCCAAATGGCCGGG | ssDNA 1 | Fig.2A/Fig.S5A |
| Target 1_gDNA_4 | P-TGCGCAAATGGCCGGG | ssDNA 1 | Fig.2A/Fig.S5A |
| Target 1_gDNA_5 | P-TGCCGAAATGGCCGGG | ssDNA 1 | Fig.2A/Fig.S5A |
| Target 1_gDNA_6 | P-TGCCCTAATGGCCGGG | ssDNA 1 | Fig.2A/Fig.S5A |
| Target 1_gDNA_7 | P-TGCCCATATGGCCGGG | ssDNA 1 | Fig.2A/Fig.S5A |
| Target 1_gDNA_8 | P-TGCCCAATTGGCCGGG | ssDNA 1 | Fig.2A/Fig.S5A |
| Target 1_gDNA_9 | P-TGCCCAAAAGGCCGGG | ssDNA 1 | Fig.2A/Fig.S5A |
| Target 1_gDNA_10 | P-TGCCCAAATCGCCGGG | ssDNA 1 | Fig.2A/Fig.S5A |
| Target 1_gDNA_11 | P-TGCCCAAATGCCCGGG | ssDNA 1 | Fig.2A/Fig.S5A |
| Target 1_gDNA_12 | P-TGCCCAAATGGGCGGG | ssDNA 1 | Fig.2A/Fig.S5A |
| Target 1_gDNA_13 | P-TGCCCAAATGGCGGGG | ssDNA 1 | Fig.2A/Fig.S5A |
| Target 1_gDNA_14 | P-TGCCCAAATGGCCCGG | ssDNA 1 | Fig.2A/Fig.S5A |
| Target 1_gDNA_15 | P-TGCCCAAATGGCCGCG | ssDNA 1 | Fig.2A/Fig.S5A |
| Target 1_gDNA_16 | P-TGCCCAAATGGCCGGC | ssDNA 1 | Fig.2A/Fig.S5A |
| Target 1_gDNA_2-3 | P-TCGCCAAATGGCCGGG | ssDNA 1 | Fig.2B/Fig.S5A |
| Target 1_gDNA_3-4 | P-TGGGCAAATGGCCGGG | ssDNA 1 | Fig.2B/Fig.S5A |
| Target 1_gDNA_4-5 | P-TGCGGAAATGGCCGGG | ssDNA 1 | Fig.2B/Fig.S5A |
| Target 1_gDNA_5-6 | P-TGCCGTAATGGCCGGG | ssDNA 1 | Fig.2B/Fig.S5A |
| Target 1_gDNA_6-7 | P-TGCCCTTATGGCCGGG | ssDNA 1 | Fig.2B/Fig.S5A |
| Target 1_gDNA_7-8 | P-TGCCCATTTGGCCGGG | ssDNA 1 | Fig.2B/Fig.S5A |
| Target 1_gDNA_8-9 | P-TGCCCAATAGGCCGGG | ssDNA 1 | Fig.2B/Fig.S5A |
| Target 1_gDNA_9-10 | P-TGCCCAAAACGCCGGG | ssDNA 1 | Fig.2B/Fig.S5A |
| Target 1_gDNA_10-11 | P-TGCCCAAATCCCCGGG | ssDNA 1 | Fig.2B/Fig.S5A |
| Target 1_gDNA_11-12 | P-TGCCCAAATGCGCGGG | ssDNA 1 | Fig.2B/Fig.S5A |
| Target 1_gDNA_12-13 | P-TGCCCAAATGGGGGGG | ssDNA 1 | Fig.2B/Fig.S5A |
| Target 1_gDNA_13-14 | P-TGCCCAAATGGCGCGG | ssDNA 1 | Fig.2B/Fig.S5A |
| Target 1_gDNA_14-15 | P-TGCCCAAATGGCCCCG | ssDNA 1 | Fig.2B/Fig.S5A |
| Target 1_gDNA_15-16 | P-TGCCCAAATGGCCGCC | ssDNA 1 | Fig.2B/Fig.S5A |
| Target 2_gDNA_0 | P-TTTGGAGCTGGTGGCG | ssDNA2 | Fig.2A/Fig.S5B |
| Target 2_gDNA_2 | P-TATGGAGCTGGTGGCG | ssDNA2 | Fig.2A/Fig.S5B |
| Target 2_gDNA_3 | P-TTAGGAGCTGGTGGCG | ssDNA2 | Fig.2A/Fig.S5B |
| Target 2_gDNA_4 | P-TTTCGAGCTGGTGGCG | ssDNA2 | Fig.2A/Fig.S5B |
| Target 2_gDNA_5 | P-TTTGCAGCTGGTGGCG | ssDNA2 | Fig.2A/Fig.S5B |
| Target 2_gDNA_6 | P-TTTGGTGCTGGTGGCG | ssDNA2 | Fig.2A/Fig.S5B |
| Target 2_gDNA_7 | P-TTTGGACCTGGTGGCG | ssDNA2 | Fig.2A/Fig.S5B |
| Target 2_gDNA_8 | P-TTTGGAGGTGGTGGCG | ssDNA2 | Fig.2A/Fig.S5B |
| Target 2_gDNA_9 | P-TTTGGAGCAGGTGGCG | ssDNA2 | Fig.2A/Fig.S5B |
| Target 2_gDNA_10 | P-TTTGGAGCTCGTGGCG | ssDNA2 | Fig.2A/Fig.S5B |
| Target 2_gDNA_11 | P-TTTGGAGCTGCTGGCG | ssDNA2 | Fig.2A/Fig.S5B |
| Target 2_gDNA_12 | P-TTTGGAGCTGGAGGCG | ssDNA2 | Fig.2A/Fig.S5B |
| Target 2_gDNA_13 | P-TTTGGAGCTGGTCGCG | ssDNA2 | Fig.2A/Fig.S5B |
| Target 2_gDNA_14 | P-TTTGGAGCTGGTGCCG | ssDNA2 | Fig.2A/Fig.S5B |
| Target 2_gDNA_15 | P-TTTGGAGCTGGTGGGG | ssDNA2 | Fig.2A/Fig.S5B |
| Target 2_gDNA_16 | P-TTTGGAGCTGGTGGCC | ssDNA2 | Fig.2A/Fig.S5B |
| Target 2_gDNA_2-3 | P-TAAGGAGCTGGTGGCG | ssDNA2 | Fig.2B/Fig.S5B |
| Target 2_gDNA_3-4 | P-TTACGAGCTGGTGGCG | ssDNA2 | Fig.2B/Fig.S5B |
| Target 2_gDNA_4-5 | P-TTTCCAGCTGGTGGCG | ssDNA2 | Fig.2B/Fig.S5B |
| Target 2_gDNA_5-6 | P-TTTGCTGCTGGTGGCG | ssDNA2 | Fig.2B/Fig.S5B |
| Target 2_gDNA_6-7 | P-TTTGGTCCTGGTGGCG | ssDNA2 | Fig.2B/Fig.S5B |
| Target 2_gDNA_7-8 | P-TTTGGACGTGGTGGCG | ssDNA2 | Fig.2B/Fig.S5B |
| Target 2_gDNA_8-9 | P-TTTGGAGGAGGTGGCG | ssDNA2 | Fig.2B/Fig.S5B |
| Target 2_gDNA_9-10 | P-TTTGGAGCACGTGGCG | ssDNA2 | Fig.2B/Fig.S5B |
| Target 2_gDNA_10-11 | P-TTTGGAGCTCCTGGCG | ssDNA2 | Fig.2B/Fig.S5B |
| Target 2_gDNA_11-12 | P-TTTGGAGCTGCAGGCG | ssDNA2 | Fig.2B/Fig.S5B |
| Target 2_gDNA_12-13 | P-TTTGGAGCTGGACGCG | ssDNA2 | Fig.2B/Fig.S5B |
| Target 2_gDNA_13-14 | P-TTTGGAGCTGGTCCCG | ssDNA2 | Fig.2B/Fig.S5B |
| Target 2_gDNA_14-15 | P-TTTGGAGCTGGTGCGG | ssDNA2 | Fig.2B/Fig.S5B |
| Target 2_gDNA_15-16 | P-TTTGGAGCTGGTGGGC | ssDNA2 | Fig.2B/Fig.S5B |
| Target 3_gDNA_2 | P-TCAAATCACTGAGCAG | ssDNA3 | Fig.2A/Fig.S5C |
| Target 3_gDNA_3 | P-TGTAATCACTGAGCAG | ssDNA3 | Fig.2A/Fig.S5C |
| Target 3_gDNA_4 | P-TGATATCACTGAGCAG | ssDNA3 | Fig.2A/Fig.S5C |
| Target 3_gDNA_5 | P-TGAATTCACTGAGCAG | ssDNA3 | Fig.2A/Fig.S5C |
| Target 3_gDNA_6 | P-TGAAAACACTGAGCAG | ssDNA3 | Fig.2A/Fig.S5C |
| Target 3_gDNA_7 | P-TGAAATGACTGAGCAG | ssDNA3 | Fig.2A/Fig.S5C |
| Target 3_gDNA_8 | P-TGAAATCTCTGAGCAG | ssDNA3 | Fig.2A/Fig.S5C |
| Target 3_gDNA_9 | P-TGAAATCAGTGAGCAG | ssDNA3 | Fig.2A/Fig.S5C |
| Target 3_gDNA_10 | P-TGAAATCACAGAGCAG | ssDNA3 | Fig.2A/Fig.S5C |
| Target 3_gDNA_11 | P-TGAAATCACTCAGCAG | ssDNA3 | Fig.2A/Fig.S5C |
| Target 3_gDNA_12 | P-TGAAATCACTGTGCAG | ssDNA3 | Fig.2A/Fig.S5C |
| Target 3_gDNA_13 | P-TGAAATCACTGACCAG | ssDNA3 | Fig.2A/Fig.S5C |
| Target 3_gDNA_14 | P-TGAAATCACTGAGGAG | ssDNA3 | Fig.2A/Fig.S5C |
| Target 3_gDNA_15 | P-TGAAATCACTGAGCTG | ssDNA3 | Fig.2A/Fig.S5C |
| Target 3_gDNA_16 | P-TGAAATCACTGAGCAC | ssDNA3 | Fig.2A/Fig.S5C |
| Target 3_gDNA_2-3 | P-TCTAATCACTGAGCAG | ssDNA3 | Fig.2B/Fig.S5C |
| Target 3_gDNA_3-4 | P-TGTTATCACTGAGCAG | ssDNA3 | Fig.2B/Fig.S5C |
| Target 3_gDNA_4-5 | P-TGATTTCACTGAGCAG | ssDNA3 | Fig.2B/Fig.S5C |
| Target 3_gDNA_5-6 | P-TGAATACACTGAGCAG | ssDNA3 | Fig.2B/Fig.S5C |
| Target 3_gDNA_6-7 | P-TGAAAAGACTGAGCAG | ssDNA3 | Fig.2B/Fig.S5C |
| Target 3_gDNA_7-8 | P-TGAAATGTCTGAGCAG | ssDNA3 | Fig.2B/Fig.S5C |
| Target 3_gDNA_8-9 | P-TGAAATCTGTGAGCAG | ssDNA3 | Fig.2B/Fig.S5C |
| Target 3_gDNA_9-10 | P-TGAAATCAGAGAGCAG | ssDNA3 | Fig.2B/Fig.S5C |
| Target 3_gDNA_10-11 | P-TGAAATCACACAGCAG | ssDNA3 | Fig.2B/Fig.S5C |
| Target 3_gDNA_11-12 | P-TGAAATCACTCTGCAG | ssDNA3 | Fig.2B/Fig.S5C |
| Target 3_gDNA_12-13 | P-TGAAATCACTGTCCAG | ssDNA3 | Fig.2B/Fig.S5C |
| Target 3_gDNA_13-14 | P-TGAAATCACTGACGAG | ssDNA3 | Fig.2B/Fig.S5C |
| Target 3_gDNA_14-15 | P-TGAAATCACTGAGGTG | ssDNA3 | Fig.2B/Fig.S5C |
| Target 3_gDNA_15-16 | P-TGAAATCACTGAGCTC | ssDNA3 | Fig.2B/Fig.S5C |
| PIK3CA primary gDNA | P-TAGTAAAAATCTCACA | PIK3CA-ssDNA | Fig. 2/ Fig. S7 |
| KRAS primary gDNA | P-TATTCGTCCACAAAAT | KRAS-ssDNA | Fig. 2 |
| NRAS primary gDNA | P-TATGTCCAACAAACAG | NRAS-ssDNA | Fig. 2 |
| EGFR primary gDNA | P-TCACTTTGCCTCCTTC | EGFR-ssDNA | Fig. 2 |
| 15nt gDNA | P-CATTTGGGCGTGCCC | ssDNA F-Q substrate | Fig. S6 |
| 20nt gDNA | P-CATTTGGGCGTGCCCCCGCA | ssDNA F-Q substrate | Fig. S6 |
| 25nt gDNA | P-CATTTGGGCGTGCCCCCGCAAGACT | ssDNA F-Q substrate | Fig. S6 |
| 30nt gDNA | P-CATTTGGGCGTGCCCCCGCAAGACTGCTAG | ssDNA F-Q substrate | Fig. S6 |
| 40nt gDNA | P-CATTTGGGCGTGCCCCCGCAAGACTGCTAGCCGAGTAGCG | ssDNA F-Q substrate | Fig. S6 |
| 50nt gDNA | P-CATTTGGGCGTGCCCCCGCAAGACTGCTAGCCGAGTAGCGTTGGGTTGCG | ssDNA F-Q substrate | Fig. S6 |
| 60nt gDNA | P-CATTTGGGCGTGCCCCCGCAAGACTGCTAGCCGAGTAGCGTTGGGTTGCGAAAGGCCTTG | ssDNA F-Q substrate | Fig. S6 |
| 70nt gDNA | P-CATTTGGGCGTGCCCCCGCAAGACTGCTAGCCGAGTAGCGTTGGGTTGCGAAAGGCCTTGTGGTACTGCC | ssDNA F-Q substrate | Fig. S6 |
| 80nt gDNA | P-CATTTGGGCGTGCCCCCGCAAGACTGCTAGCCGAGTAGCGTTGGGTTGCGAAAGGCCTTGTGGTACTGCCTGATAGGGCG | ssDNA F-Q substrate | Fig. S6 |
| HPV6-gDNA | P-TCTACATCTTGCACAT | HPV6 containing plasmid | Fig. 3 |
| HPV11-gDNA | P-TCTAAATCTGGTACAT | HPV11 containing plasmid | Fig. 3 |
| HPV16-gDNA | P-TCATGTCGTTGGTACT | HPV16 containing plasmid | Fig. 3 |
| HPV18-gDNA | P-TCATGTCTGCTATACT | HPV18 containing plasmid | Fig. 3 |

**Table S3. ssDNA targets used in this study.**

| Name | Sequence | Presence |
| --- | --- | --- |
| Primary ssDNA | GGTCCTTTCTTGGATAAACCCACTCTATGCCCGGCCATTTGGGCGTGCCCCCGCAAGACT | Fig.1C/Fig.S3B |
| Secondary ssDNA | AGTCTTGCGGGGGCACGCCCAAATGGCCGGGCATAGAGTGGGTTTATCCAAGAAAGGACC | Fig.1C |
| ssDNA1 | AGTCTTGCGGGGGCACGCCCAAATGGCCGGGCATAGAGTGGGTTTATCCAAGAAAGGACC | Fig.2A/Fig.S5A |
| ssDNA2 | TAGCTGTATCGTCAAGGCACTCTTGCCTACGCCACCAGCTCCAACTACCACAAGTTTATA | Fig.2A/Fig.S5A |
| ssDNA3 | CTCCATAGAAAATCTTTCTCCTGCTCAGTGATTTCAGAGAGAGGATCTCGTGTAGAAATT | Fig.2A/Fig.S5A |
| PIK3CA-ssDNA | AAATATGAACAATATTTGGATAACTTGCTTGTGAGATTTTTACTGAAGAAAGCATTGACT | Fig. 2/ Fig. S7 |
| KRAS-ssDNA | TGCCTTGACGATACAGCTAATTCAGAATCATTTTGTGGACGAATATGATCCAACAATAGA | Fig. 2 |
| NRAS-ssDNA | AGAAAACAAGTGGTTATAGATGGTGAAACCTGTTTGTTGGACATACTGGATACAGCTGGA | Fig. 2 |
| EGFR-ssDNA | TGGGTGCGGAAGAGAAAGAATACCATGCAGAAGGAGGCAAAGTGCCTATCAAGTGGATGG | Fig. 2 |

**Table S4. primers used in this study.**

| Name | Target | Forward primer sequence | Presence |
| --- | --- | --- | --- |
| JHF 600bp-FW | JHF containing plasmid | CCGGTGAGTACACCGGAATTGC | Fig. 1 |
| JHF 600bp-RV | JHF containing plasmid | GCAGTCTTGCGGGGGC | Fig. 1 |
| JHF 95bp-FW | JHF containing plasmid | CGACACTCCGCCATGAATCAC | Fig. 1 |
| JHF 95bp-RV | JHF containing plasmid | CAGCCGAGTCCCTCATTCCC | Fig. 1 |
| HPV-universal -FW | HPV containing plasmids | CYACWCGCAGTACMAAYWTRWCAHTATGTGC | Fig. 3 |
| HPV-universal -RV | HPV containing plasmids | TTGAAAAATAAAYTGYAAATCAWAYTCYTC | Fig. 3 |

**Table S5. Cleavage reporters used in this study.**

| Name | Sequence | Fluoro-  phore | Presence |
| --- | --- | --- | --- |
| PIK3CA reporter | /56-FAM/TAGTCAATGCTTTCTTCAGTAAAAATCTCA/3BHQ1-FQ/ | FAM | Fig. 2/ Fig. S7 |
| KRAS reporter | /5-JOE/CTCTATTGTTGGATCATATTCGTCCACAAA/3BHQ1-FQ/ | JOE | Fig. 2 |
| NRAS reporter | /5-NED/GTCCAGCTGTATCCAGTATGTCCAACAAAC/3BHQ2-FQ/ | NED | Fig. 2 |
| EGFR reporter | /5-ROX/GCCATCCACTTGATAGGCACTTTGCCTCCT/3BHQ2-FQ/ | ROX | Fig. 2 |
| ssDNA F-Q substrate | /56-FAM/CTCGGCTAGCAGTCTTGCGGGGGCACGCCCAAATGGCCGG/3BHQ1-FQ/ | FAM | Fig. S6/ Fig. S8 |
| 20nt reporter | /56-FAM/GGGGCACGCCCAAATGGCCG/3BHQ1-FQ/ | FAM | Fig. S8 |
| 22nt reporter | /56-FAM/CGGGGGCACGCCCAAATGGCCG/3BHQ1-FQ/ | FAM | Fig. S8 |
| 24nt reporter | /56-FAM/TGCGGGGGCACGCCCAAATGGCCG/3BHQ1-FQ/ | FAM | Fig. S8 |
| 26nt reporter | /56-FAM/CTTGCGGGGGCACGCCCAAATGGCCG/3BHQ1-FQ/ | FAM | Fig. S8 |
| 30nt reporter | /56-FAM/CAGTCTTGCGGGGGCACGCCCAAATGGCCGCATTTGGGCGTGCCCC/3BHQ1-FQ/ | FAM | Fig. S6/  Fig. S8 |
| HPV6 reporter | /5-NED/ATTATGTGCATCCGTAACTACATCTTCCAC /3BHQ2-FQ/ | NED | Fig. 3 |
| HPV11 reporter | /5-ROX/ACTATGTGCATCTGTGTCTAAATCTGCTAC /3BHQ2-FQ/ | ROX | Fig. 3 |
| HPV16 reporter | /56-FAM/TAAATCATATTCCTCCCCATGTCGTAGGTA /3BHQ1-FQ/ | FAM | Fig. 3 |
| HPV18 reporter | /5-JOE/CAAATCATATTCCTCAACATGTCTGCTATA /3BHQ1-FQ/ | JOE | Fig. 3 |

**Table S6. Plasmids used in this study.**

| Plasmid name | Description |
| --- | --- |
| *Pf*Ago-plasmid | pET28a-*Pf*Ago |
| PIK3CA-plasmid | pET28a-PIK3CA |
| JHF1-plasmid | pET28a-JHF1 |
| HPV-type6- plasmid | pET28a-HPV-type6-Gene 1-Loop 5 |
| HPV-type11- plasmid | pET28a-HPV-type11-Gene 1-Loop 5 |
| HPV-type16- plasmid | pET28a-HPV-type16-Gene 1-Loop 5 |
| HPV-type18- plasmid | pET28a-HPV-type18-Gene 1-Loop 5 |

**Table S7. Nucleic acid for EMAS used in this study.**

| Name | Sequence | Fluorophore | Presence |
| --- | --- | --- | --- |
| EMSA-gDNA | P-TATTCGTCCACAAAATGATTCTGAATTAGC | / | Fig. S2 |
| EMSA-gDNA-FAM | P-TATTCGTCCACAAAATGATTCTGAATTAGC-6-FAM | FAM | Fig. S2 |
